# Observation of histone nuclear import in living cells: implications in the processing of newly synthesised H3.1 & H4

**DOI:** 10.1101/111096

**Authors:** Michael James Smith, Andrew James Bowman

## Abstract

**Highlights:**

- Small-molecule-gated tether-and-release system for rapid pulse-chase of nuclear proteins
- Tracking nuclear import of histone H3.1 and H4 and their incorporation at sites of active replication
- Tethered H3.1 and H4 are monomeric and do not associate with ASF1, NASP, RbAp46 or HAT1 in the cytosol
- Importin-β proteins as cytosolic binders of monomeric histones

**Summary:** We present here a cytosolic tether-and-release system to study the import and dynamics of newly synthesised nuclear proteins. Release is gated by rapamycin-induced recruitment and activation of a viral protease, with cleavage of a peptide linker releasing the tethered cargo. We use this system to investigate nucleo-cytoplasmic divisions in the histone H3.1 & H4 deposition pathway, revealing that, contrary to previous analyses, H3.1 and H4 are predominantly monomeric in the cytosol, and only associate with the core histone chaperoning machinery after translocation to the nucleus. Whilst we do not detect interaction with known H3-H4 chaperones in the cytosol we do detect interaction with a number of importin-β proteins, that may serve a dual import and chaperoning function, preventing aggregation of histones until they are handed-off to the core histone chaperoning machinery in the nucleus.

## Introduction

Chromatin is the cellular template with which factors involved in genomic transactions have to contend. As the central component of chromatin, understanding how histones are deposited onto DNA, how this process is coordinated with DNA replication and nucleosome turnover, and how parental histones are segregated to daughter strands has implications in many areas of biology. Each cell division requires the doubling of both DNA and histone content, with half of the histones being of parental origin and half being newly synthesized. Whilst much effort has gone into studying the dynamics of recycled parental histones (Alabert et al., 2015;Jackson, 1987, 1990;Katan-Khaykovich and Struhl, 2011;Prior et al., 1980;Radman-Livaja et al., 2011), less is known about the program for newly synthesized histone incorporation. As they form the stable core of the nucleosome and are the substrate for the majority of post-translational marks, histones H3 and H4 are often at the forefront of these investigations.

Unlike recycled histones, newly synthesized histones H3 and H4 must pass through the cytosol before they are incorporated into chromatin. Biochemical isolation of H3.1 (the replication dependent H3 variant) containing complexes suggested it folds with H4 soon after synthesis, interacting with a number of histone chaperones to form a cytosolic chaperoning network that coordinates nuclear import (Alvarez et al., 2011;Ask et al., 2012;Campos et al., 2010;Mosammaparast et al., 2002). Key cytosolic events in the proposed pathway include H3.1 and H4 forming a heterodimer in the cytosol and interacting with NASP, ASF1, HAT1 and RbAp46 (Alvarez et al., 2011;Campos et al., 2010;Mosammaparast et al., 2002), HAT1 modification of H4 K5 and K12 by acetylation [reviewed in (Parthun, 2011)] (Alvarez et al., 2011), modification H3 K9 by methylation (Pinheiro et al., 2012;Rivera et al., 2015) and association with the Importin-β protein IPO4 (Ask et al., 2012;Blackwell et al., 2007;Campos et al., 2010;Gurard-Levin et al., 2014;Hammond et al., 2017;Keck and Pemberton, 2012;Mosammaparast et al., 2002). Indeed, previous to the identification of cytosolic histone chaperones, a number of Importin-β proteins were suggested to act as cytosolic chaperones for the core histones (Jakel et al., 2002).

Nuclear import and incorporation of histones into chromatin occurs very rapidly (Ruiz-Carrillo et al., 1975), thus following such events in living cells remains challenging. Import rates most likely exceed the folding and maturation kinetics of fluorescent proteins, making the process difficult to study by FRAP, FLIP, photoactivation, or their derivative techniques (Ishikawa-Ankerhold et al., 2012;Lukyanov et al., 2005;Reits and Neefjes, 2001). Similar difficulties arise with self-labelling domains such as the SNAP-tag, requiring minutes to hours for quenching, pulsing and labelling steps (Clement et al., 2016;Crivat and Taraska, 2012;Jansen et al., 2007;Juillerat et al., 2003). Metabolic incorporation of radioactive amino acids or functional amino acid derivatives (Deal et al., 2010;Dieterich et al., 2007;Lang and Chin, 2014) affords immediate labelling for biochemical analysis, but presents challenges for imaging proteins due to the requirement for derivatisation of the incorporated functional groups (Lang and Chin, 2014). Thus, many of the current ideas about the nucleo-cytoplasmic chaperoning of histones remain to be tested in a cellular setting.

In an attempt to address this, and potentially gain new information regarding the histone import and deposition pathway, we have developed an approach termed RAPID-release (<u>R</u>apamycin <u>A</u>ctivated <u>P</u>rotease through <u>I</u>nduced <u>D</u>imerisation and <u>release</u> of tethered cargo) that allows observation of dynamic cellular events in real-time in living cells. In this approach we circumvent the requirement for immediate labelling of newly synthesized histones by first capturing them on the cytosolic side of the outer mitochondrial membrane (OMM). The quiescent histones are then released by concomitant recruitment and activation of a site-specific, viral protease through addition of the small molecule rapamycin (Stein and Alexandrov, 2014). The histones can be followed by fusion of a fluorescent protein, allowing visualization of nuclear import and incorporation at replication domains in real time. We apply this approach to investigate the early maturation and import of H3.1 and H4.

## Results

### The RAPID-release technique allows observation of histone nuclear import

RAPID-release allows the accumulation of a target protein in the cytosol by tethering it to the OMM (Figure 1A). This permits fluorescent fusion proteins to fold and mature, generating a fluorescence pool in the cytosol for pulse-chase analysis. The release of the tethered target is triggered by addition of rapamycin, which concomitantly recruits and activates an auto-inhibited TVMV protease (Stein and Alexandrov, 2014).

**Figure 1.**
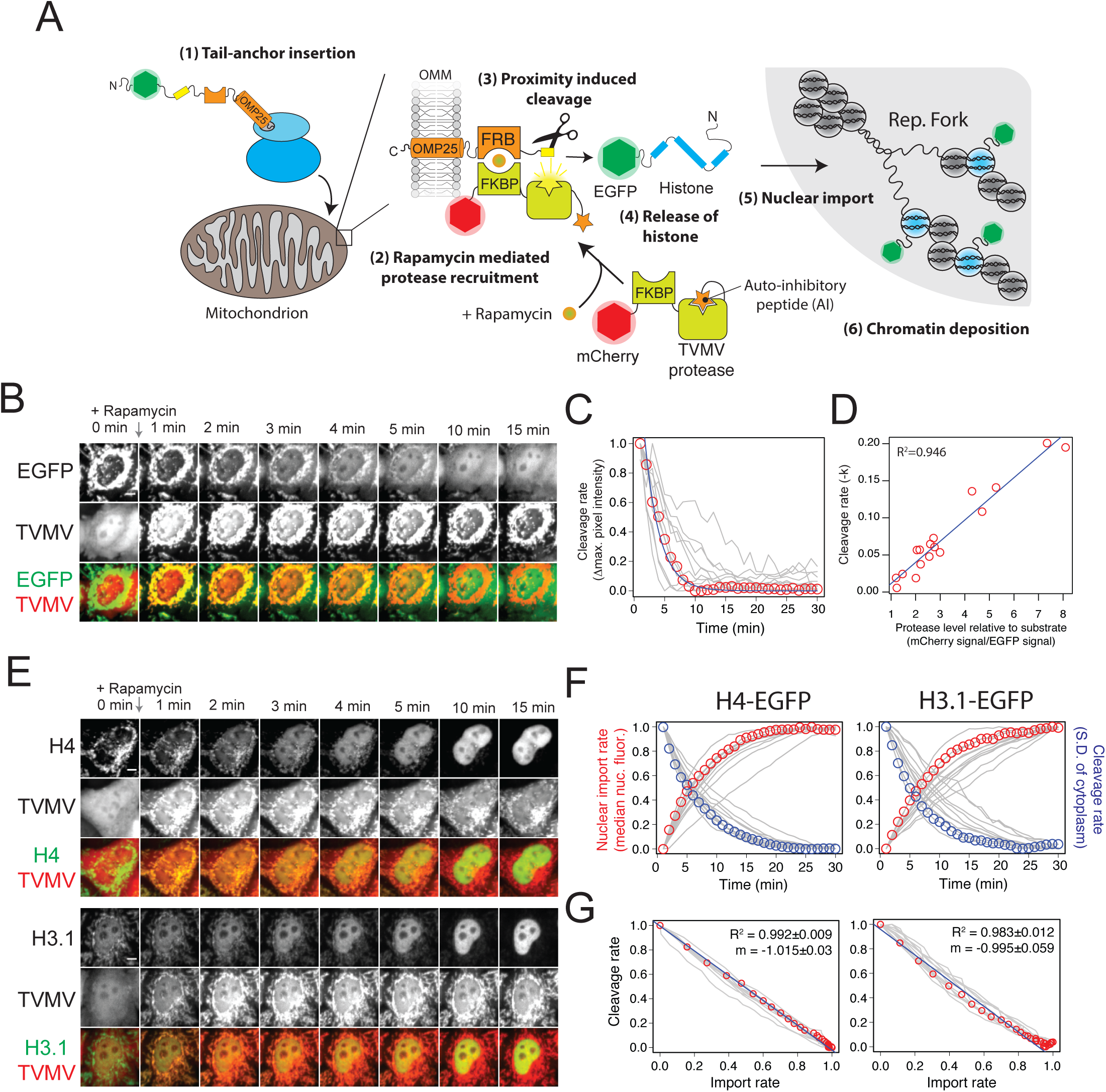
Observation of histone nuclear import in living cells. (A) Schematic representation of the RAPID-release technique. Addition of rapamycin results in recruitment and activation of an auto-inhibited TVMV protease, leading to the release of EGFP-labelled histones from their mitochondrial tether. (B) A representative cell showing release of tethered EGFP. Scale bar represents 5 μm. (C) Quantification of EGFP release. Single cells are represented as grey traces. The cell displayed in B is highlighted with red circles. The blue line represents a fit to an exponential decay model. (D) Rate constants from exponential decay functions of the traces shown in C plotted against the expression level of protease relative to EGFP substrate. The blue line represents a linear regression model with the R value shown. (E) Representative cells showing release of H4-EGFP and H3.1-EGFP. Scale bar represents 5 μm. (F) Nuclear import rate and cleavage rate of H3.1-EGFP and H4-EGFP. Single cells are represented as grey traces. Traces from cells displayed in E are highlighted with circles. (G) Import rate plotted against cleavage rate. Values represent those shown in F, with the cells shown in E highlighted with red circles. Linear regression models are shown with blue lines. The mean R and slope (m) are shown for the 11 cells quantified in F, +/- relates to standard deviation.

To assess the feasibility of the approach we fused EGFP to the FKBP12-rapamycin-binding (FRB) domain of mTOR, followed by the mitochondrial tail-anchoring sequence of OMP25 (Horie et al., 2002), with two TVMV cleavage sites separating the EGFP and FRB-OMP25 domains (EGFP^TVMVx2^-FRB-OMP25) (Figure 1A). The TVMV protease containing a C-terminal Auto-Inhibitory (AI) peptide and an N-terminal FK506-binding domain (FKBP12) (Stein and Alexandrov, 2014) was fused to the C-terminus of mCherry, creating the construct mCherry-FKBP12-TVMV-AI (Figure 1A).

Initially we used EGFP without a fusion partner to test the feasibility of the approach. HeLa cells were chosen as this cell line has been used most frequently in the biochemical characterisation of the H3.1-H4 chaperoning pathway (Campos et al., 2010;Groth et al., 2005;Tagami et al., 2004). Addition of rapamycin to HeLa cells co-transfected with EGFP^TVMVx2^-FRB-OMP25/mCherry-FKBP12-TVMV-AI resulted in the recruitment of mCherry-FKBP12-TVMV-AI to mitochondria and release of the EGFP cargo from its tether (Figure 1B). Analysis of the cleavage (as the change in maximum pixel intensity of the cytoplasm) revealed a fit to an exponential decay model (Figure 1C). Plotting the rate constant of fit against the relative expression level of the protease (measured as the ratio of mCherry:EGFP signal) for each cell revealed a strong positive correlation, suggesting the cleavage rate is dependent on the level of protease (Figure 1D). At the highest ratios of protease to substrate a half maximal cleavage of 2.5 minutes was achieved. Removal of the two TVMV cleavage sites from EGFP inhibited cleavage, whilst removal of the AI peptide from the TVMV protease resulted in constitutive activity without recruitment (Figure S1A & B), demonstrating the specificity of the protease and the importance of the AI peptide fusion, respectively.

Next we used the RAPID-release system to observe the dynamics of histone H3.1 and H4 upon release from the OMM. In this instance, tail-anchoring, in contrast to N-terminal anchoring (Suzuki et al., 2002), permitted C-terminal tagging of histones, allowing us to avoid N-terminal fusions that have previously been shown to affect chromatin incorporation dynamics (Kimura and Cook, 2001). Transient transfection of H3.1 or H4 fused to the N-terminus of the EGFP^TVMVx2^-FRB-OMP25 construct did not observably affect mitochondrial or cellular morphology. However, a small amount of background nuclear fluorescence was observed (Figure 1E, Movie S1 & S2). As tail-anchor insertion is a post-translational event (Borgese and Fasana, 2011;Krumpe et al., 2012;Mariappan et al., 2011;Okreglak and Walter, 2014;Shao and Hegde, 2011), and as nuclear fluorescence was also observed in the absence of TVMV protease, it is likely that the background relates to histone import before OMM insertion can occur. Release of H3.1/H4-EGFP from the cytosolic tether resulted in rapid nuclear localisation (measured as the median nuclear fluorescence) (Figure 1E) at a rate mirrored by the kinetics of cleavage (measured as the S.D of the cytoplasm) (Figure 1F & G). Standard deviation of the cytoplasm was used instead of maximum pixel as it was less affected by sub-cellular partitioning. Plotting the nuclear import rate against the cleavage rate resulted in a strong fit to a linear model (Figure 1G), revealing nuclear import of histones occurs at a rate greater than proteolytic cleavage and in excess of our sampling rate. Confirming this, the modal value of the partitioned cytoplasmic signal, representing the portion of the cytoplasm outside of the mitochondrial network, did not increase over the cleavage period (Figure S1C) as it did for freely diffusing EGFP.

In summary, the RAPID-release technique allows observation of histone nuclear import in living cells, and provides a pulse-labelling strategy with significantly improved kinetics compared to currently available techniques.

### Released histones incorporate at actively replicating domains

In a subset of asynchronously dividing cells we observed foci forming in the nucleus after histone release (Figure 2A). Mammalian genomes are organized into topological domains (TADs), which are also the stable units of replication, or Replication Domains (RDs) (Pope et al., 2014;Rivera-Mulia and Gilbert, 2016). To determine if these foci represent histone incorporation at replicating domains we co-transfected H3.1-EGFP^TVMVx2^-FRB-OMP25 fusions with a PCNA-V_H_H-TagRFP chromobody, a marker of active replication (Burgess et al., 2012). FKPB-TVMV-AI was also co-transfected, but as a TagBFP fusion to allow imaging of PCNA. The TagBFP channel was used to identify expressing cells, but was not imaged to minimise bleaching of the other fluorophores. Fixing and imaging cells 30 minutes after rapamycin addition showed that nuclei positive for PCNA foci were also positive for H3.1-EGFP foci, whereas nuclei negative for PCNA foci were also negative for H3.1-EGFP foci (Figure 2B). Cells absent of PCNA foci, and by deduction not replicating their genomes, demonstrated a nucleolar enrichment of H3.1-EGFP, which has previously been shown to be an artefact of excess soluble histones (Musinova et al., 2011;Safina et al., 2017).

**Figure 2.**
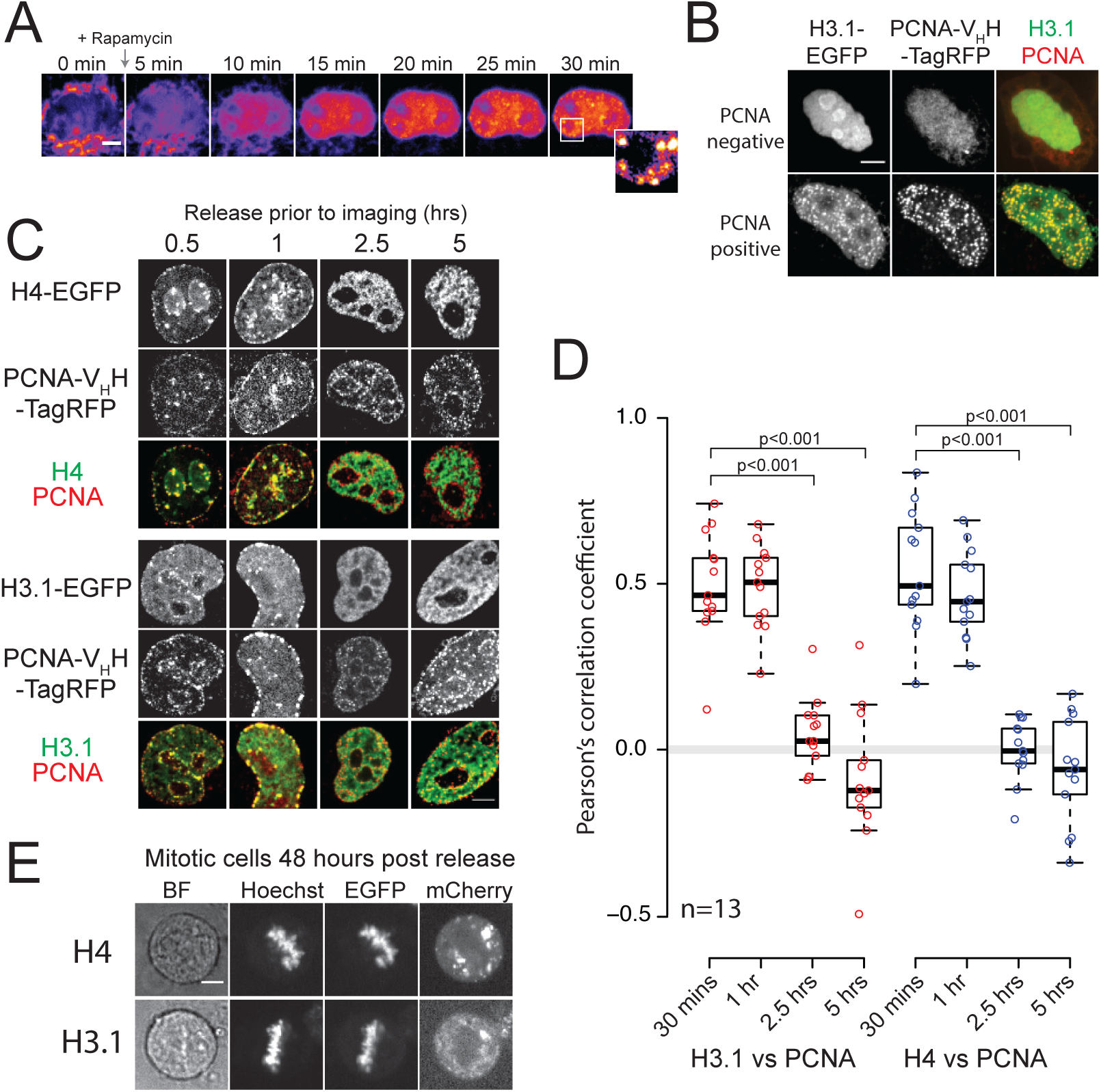
Incorporation of released histones into chromatin at sites of active replication. (A) Accumulation of nuclear H3.1-EGFP and the formation of distinct foci. Scale bar represents 5 μm. (B) Representative cells showing two different subpopulations of released H3.1-EGFP staining and their colocalization with PCNA positive foci. PCNA was detected with a PCNA-V_H_H-TagRFP chromobody. Scale bar represents 5 μm. (C) Representative cells showing points from a time course of H3.1-EGFP and H4-EGFP release. Scale bar represents 5 μm. (D) Colocalization of PCNA and H3.1/H4-EGFP signals at various time points after release. A Wilcoxon rank sum test was carried out for each time point versus the 30 min time point, with those scoring a p-value of <0.001 indicated by brackets. (E) Released H3.1/H4-EGFP colocalize with mitotic chromosomes 48 hours post release. Scale bar represents 5 μm.

To analyse the incorporation dynamics more quantitatively we released histones at four time points prior to fixation and co-localisation analysis. Cells in mid/late S-phase were chosen, identified from their peripheral PCNA staining pattern (Burgess et al., 2012), as these cells would have already entered S-phase when the histones were released from their cytosolic tether. Within 30 minutes formation of foci occurred in the EGFP channel that colocalised to replication domains marked by the PCNA chromobody (Figure 2C and D). Colocalisation continued for one hour, then decreased dramatically at 2.5 hours, with a slight anti-correlation seen at five hours post-release (Figure 2C and D). We interpret this as the pulse of released histones entering the soluble pool and being incorporated at actively replicating domains, up until the fluorescent pool of histones is depleted and replication moves on to neighbouring domains.

To further verify that released histones are incorporated into chromatin we cultured cells 48 hours after rapamycin addition before imaging those in mitosis. We found that the H3.1- /H4-EGFP signal localised to the condensed, mitotic chromosomes, providing further evidence that released H3.1-/H4-EGFP are deposited into chromatin and remain stably associated with chromatin through multiple cell divisions (Figure 2E). Together these experiments demonstrate that released H3.1-/H4-EGFP enter the histone chaperoning pathway and are incorporated into chromatin in a similar fashion to endogenous histones, and validates the RAPID-release system as a method for investigating chromatin assembly and the histone chaperoning pathway.

### Tethered, cytosolic H3.1 and H4 are monomeric and do not associate with endogenous NASP, ASF1A, RbAp46 or HAT1

H3.1 and H4 exist as an obligate heterodimer in chromatin and are also found as a dimer when bound to a number of histone chaperoning proteins, such as ASF1A/B, s/tNASP, RbAp46, HAT1 and the CAF1 complex (Tagami et al., 2004). However, the two histones are synthesised separately and must fold at a point prior to entry into the chromatin deposition pathway. As an H3.1-H4 heterodimer bound to ASF1, NASP, RbAp46 and HAT1 has previously been isolated from HeLa cytoplasmic extracts (Ask et al., 2012;Campos et al., 2010;Cook et al., 2011;Drane et al., 2010;Groth et al., 2007;Tagami et al., 2004) we wondered whether H3.1 and H4 exist in this state when coupled to their mitochondrial tether in our RAPID-release system. We reasoned that if H3.1 and H4 fold in the cytoplasm, we would expect to see the endogenous partner of each histone being enriched on the mitochondrial network. Similarly, endogenous histone chaperones that interact with H3.1 and H4 should also be enriched at the mitochondrial network (Figure 3A). To test this we carried out immunofluorescence to probe for the co-occurrence of endogenous histone binding partners (Figure 3B & C). Profiles were measured across the mitochondrial network for each staining and the correlation between the EGFP and immunofluorescent signals quantified (Figure 3C & S2A, see material and methods for details of correlation analysis). Interestingly, whilst we could detect the tethered histones, we could not detect enrichment of their orthologous binding partners, nor could we detect any of the histone chaperones previously proposed to be cytosolic (Figure 3C & S2A). Indeed, contrary to previous biochemical analysis, all of the chaperones appeared to be exclusively nuclear in their staining pattern (Figure S2B).

**Figure 3.**
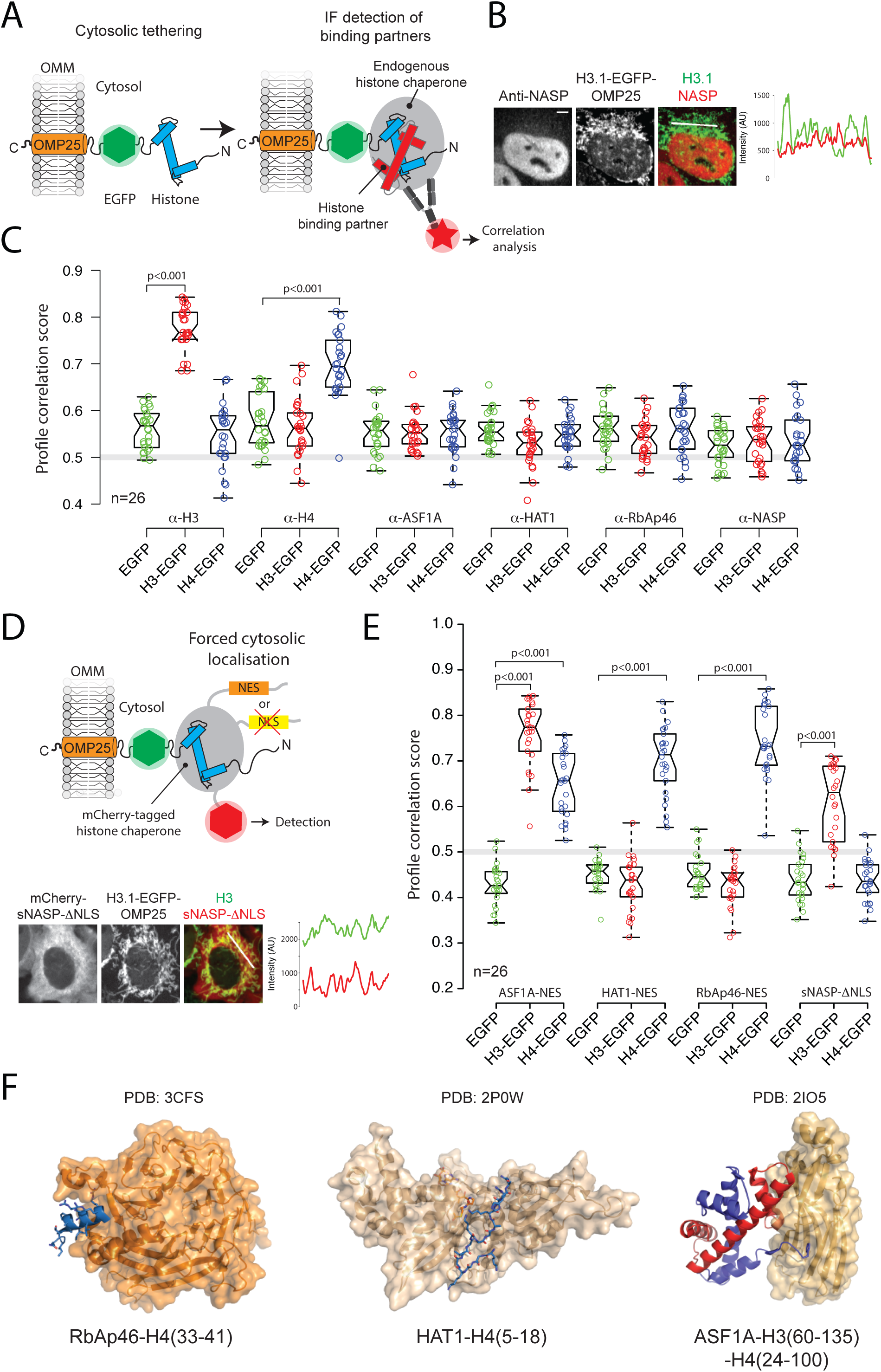
Interaction profiling of tethered cytosolic histones. (A) Schematic representation of the fluorscence-2-hybrid approach for analysing interaction with endogenous proteins. (B) Representative cell expressing H3.1 tethered to mitochondria and stained with a NASP antibody. The 1D profile corresponds to the white line shown in the merged panel. Scale bar represents 5 μm. (C) Profile correlation scores between tethered H3.1/H4-EGFP and endogenous histone counterparts or known histone chaperones. The grey line represents a zero correlation level. P-values of <0.001 from a Wilcoxon rank sum test versus the EGFP alone control are indicated by brackets. Whiskers extend to 1.5 times the IQR, notches extend to 1.58 IQR divided by the square root of n. (D) Schematic representation of a fluorescence-2-hybrid assay using forced cytosolic chaperones expressed from trans-genes with representative cell and profile shown. Scale bar represents 5 μm. (E) Profile correlation scores between tethered H3.1/H4-EGFP and cytosolically forced chaperones. P-values of <0.001 from a Wilcoxon rank sum test versus the EGFP control are indicated by brackets. The grey line represents a zero correlation level. Whiskers extend to 1.5 times the IQR, notches extend to 1.58 IQR divided by the square root of n. (F) Crystal structures of RbAp46-H4, HAT1-H4 and ASF1A-H3-H4 complexes. H3 is shown in red, H4 is shown in blue.

To determine if lack of binding was due to the nuclear partitioning of histone chaperones, rather than tethered histones adopting an unfavourable conformation that prevents binding, we expressed forced cytosolic chaperones that were mCherry-tagged. To achieve cytosolic localisation chaperones were either mutated in their nuclear localisation sequence (ΔNLS), where a defined NLS existed (as for NASP (Kleinschmidt and Seiter, 1988; O'Rand et al., 1992)), or engineered with a strong nuclear export signal (NES) (Henderson and Eleftheriou, 2000) where no known NLS was present (as for RbAp46, HAT1, ASF1A) (Figure 3D). This effectively drove the cytosolic location of all histone chaperones tested (Figure S2C). The rational behind the experiment was as follows. As RbAp46 and HAT1 bind to H4 epitopes within the H3.1-H4 heterodimer (Murzina et al., 2008;Song et al., 2008) (Figure 3F), if tethered histones were folded with their endogenous counterpart we would expect to see the recruitment of RbAp46 and HAT1 to both tethered H3.1 and tethered H4, whereas, if histones were monomeric we would expect to see recruitment to tethered H4, but not H3.1. Conversely, as sNASP interacts directly with H3 as a monomer and as an H3.1-H4 heterodimer (Bowman et al., 2017;Bowman et al., 2016), we would expect to see recruitment to both tethered H3.1 & H4 if a heterodimer was present, but only to tethered H3.1 if the histones were monomeric. As ASF1 contacts both H3 and H4 through independent binding sites (English et al., 2005;Natsume et al., 2007), we may expect to see recruitment to both independent of the histone’s oligomeric status. Remarkably, forced cytosolic RbAp46-NES and HAT1-NES both localized to tethered H4 but not H3.1, whereas sNASP-ΔNLS localised to tethered H3.1 but not H4 (Figure 3E & S2C). As expected, due to having distinct binding sites for each histone (Figure 3E), ASF1A interacted with both H3.1, and, to a lesser extent H4.

Taking into consideration the inability to detect the endogenous histone counterparts of tethered H3.1 and H4, and the binding profiles of the forced cytosolic chaperones, our results suggest that the tethered histones reside in their monomeric form, and do not associate with endogenous ASF1A, s/tNASP, RbAp46 or HAT1, which appear to be predominantly nuclear in location.

### Nuclear-cytoplasmic partitioning of histone chaperones is rapidly lost during biochemical fractionation

To reconcile our findings with current knowledge regarding histone chaperoning across the nucleo-cytoplasmic divide, we further investigated the discrepancies seen between biochemical fractionation and immunofluorescence. Confirming previous reports (Campos et al., 2010;Groth et al., 2007;Groth et al., 2005), fractionation of cytosolic and nuclear compartments using a standard NP-40 lysis protocol (Suzuki et al., 2010) revealed a cytoplasmic location for s/tNASP, HAT1, RbAp46 and ASF1A, whereas CAF1p60 was both in the cytoplasmic and nuclear fractions (Figure 4A). Histone H3 was almost entirely in the nucleus, with only a trace amount in the cytoplasm. Tubulin was entirely cytosolic, whereas Lamin A/C was entirely nuclear. Puzzlingly, when the same antibodies used in western blot detection were used in immunofluorescence the histone chaperones were found to be entirely nuclear (Figure 4B & C). Pre-blocking the antibodies with their immunogen reduced nuclear staining to background levels, but did not affect the background cytoplasmic fluorescence (Figure S3A). As histone expression is cyclical, peaking in S-phase, we also tested if the cell cycle had an effect on chaperone localization, but found no difference in nuclear staining between those cells in S-phase and those cells out of S-phase (cells in S-phase were determined by the presence of PCNA) (Figure 4B). The CAF1p60 antibody was the only antibody to show colocalisation with PCNA foci, supporting the presence of the CAF1 complex at the replication fork (Ben-Shahar et al., 2009;Krude, 1995;Shibahara and Stillman, 1999). We also tested the localisation of EGFP-fusions for ASF1A, sNASP, RbAp46, HAT1 and CAF1p60 and found them all to be nuclear in their localisation, both during and outside of S-phase (Figure S3B).

**Figure 4.**
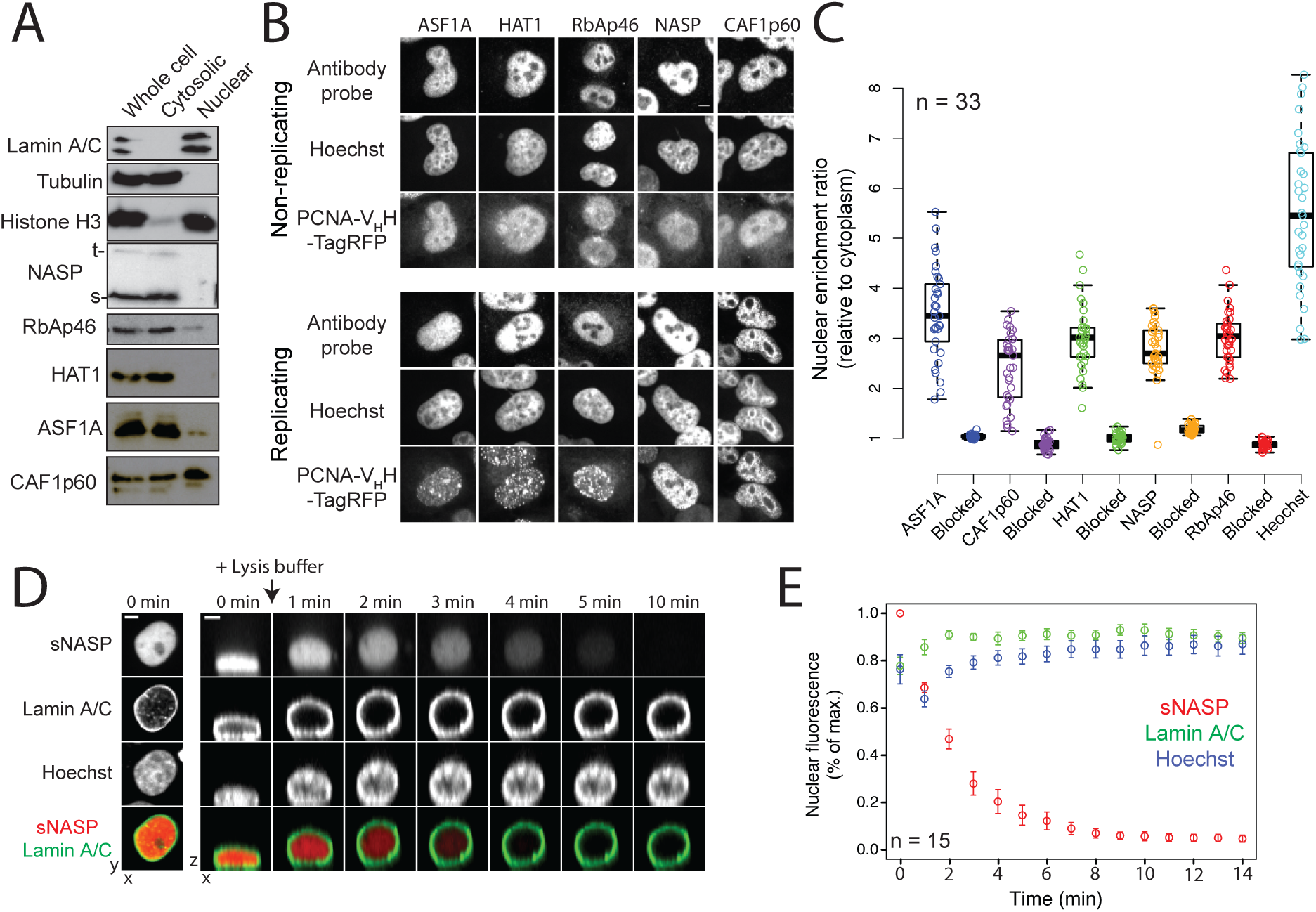
Cellular localisation of core H3.1 & H4 chaperoning components. (A) Fractionation of nuclear and cytosolic compartments using a standard NP-40 lysis protocol and immunoblotting with chaperone specific antibodies. (B) Immunofluorescence of histone chaperones in fixed cells using the same antibodies used in A. Cells are segregated into those undergoing replication and those that are not. Scale bar represents 5 μm. (C) Quantification of nuclear localisation shown in B. (D) Real-time imaging of cells undergoing biochemical fractionation. sNASP is tagged with mCherry, whilst LamnA/C is tagged with EGFP. The left column of panels represents a maximum intensity projection. The time course on the right represents a plane in the z dimension reconstructed from 20 z-stacks penetrating 20 μm into the medium. Scale bars represent 5 μm. (E) Quantification of nuclear leakage shown in D. Z-stacks were flattened using a maximum intensity projection with the nuclear fluorescent signal over time plotted as a percentage of maximum (normalised). Data points represent the mean of 15 measurements with error bars representing the s. e. m..

As the plasma membrane is ruptured during cellular fractionation it is likely that components that maintain the nucleo-cytoplasmic division, including the RanGTP-RanGDP gradient, are lost, allowing free diffusion of anything small enough to fit through the nuclear pore complex. To test how this affects the real-time localisation of histone chaperones we imaged mCherry-sNASP whilst the cell medium was replaced with lysis buffer (Figure 4D). EGFP-fused Lamin A/C was used to visualise the nuclear periphery and served as a control for nuclear integrity (Figure 4A), whilst Hoechst was used to stain the genomic DNA. Addition of lysis buffer to the cells caused the nuclei, which are normally flattened against the well of the culture dish, to puff-up (Figure 4D), due to the loss of constraining cytoskeletal forces. Crucially, the cells remained in close proximity to their pre-lysis position allowing analysis of diffusion kinetics after lysis. To follow sNASP localisation z-stacks penetrating 20 μm into the culture medium, covering the puffed-up nuclei, were taken every minute. Interestingly, whilst Lamin A/C remained at the nuclear periphery throughout the time course, sNASP rapidly diffused out of the nucleus, falling below the level of detection within 5 minutes (Figure 4D and C, Movie S3). Similar behaviour was seen for hypotonic lysis in which cells were monitored whilst being exposed to water (Figure S3C). It should be noted that previous studies using Xenopus oocytes have recorded similar findings looking at total nuclear protein levels (Paine et al., 1983;Paine et al., 1992). In summary, we propose that the discrepancy between biochemical observations and *in situ* observations can be explained in terms of the rapid diffusion of soluble nuclear components during fractionation of the cytoplasmic and nuclear compartments, and that the core histone chaperoning proteins NASP, ASF1A, RbAp46 and HAT1 are nuclear in their localisation both inside and outside of S-phase.

### The Importin-β nuclear receptors IPO4 and KPNB1 are cytosolic histone chaperones

Previous reports suggested importins may play a dual role in both chaperoning basic nuclear proteins and acting as receptors in delivering them to the nucleus (Jakel et al., 2002). Indeed, human IPO4 (also known as Imp4, Imp4b & RanBP4) has been isolated bound to H3.1 and H4 in complex with ASF1 from HeLa cell extracts (Ask et al., 2012;Campos et al., 2010;Jasencakova et al., 2010). To further investigate whether importins interact with our cytosolically tethered histones we screened three importin-β family members - IPO4, KPNB1 (IMB1, PTAC97, NTF97) and IPO11 (Imp11, RanBP11) for co-localisation to the mitochondrial network (Figure 5A & B). Based on recent phylogenetic analysis of the Importin-β family (O'Reilly et al., 2011) IPO4 and KPNB1 were chosen as representatives of two closely related branches of the import exclusive importin-β superfamily, whereas IPO11 was chosen as a member of a more distantly related branch (Figure 5C). We screened a number of antibodies to these Importins, but could not find any that were suitable for immunofluorescence, and soassessed interaction through colocalisation of mCherry-importin fusions. Remarkably, we found that IPO4 and KPNB1 interacted with tethered H4, whereas IPO4, but not KPNB1, interacted with tethered H3.1 (Figure 5A and B). IPO11 didn't interact with either tethered H3.1 or H4, and in contrast to IPO4 and KPNB1, was predominantly located in the nucleus (Figure 5A). mCherry on it own served as a control for non-specific binding. Interestingly, a recent crystal structure has identified the Importin-β protein TNPO1 (KPNB2, MIP1), closely related to KPNB1 (O'Reilly et al., 2011), as interacting with the H3 tail (Figure S4) (Soniat and Chook, 2016).

**Figure 5.**
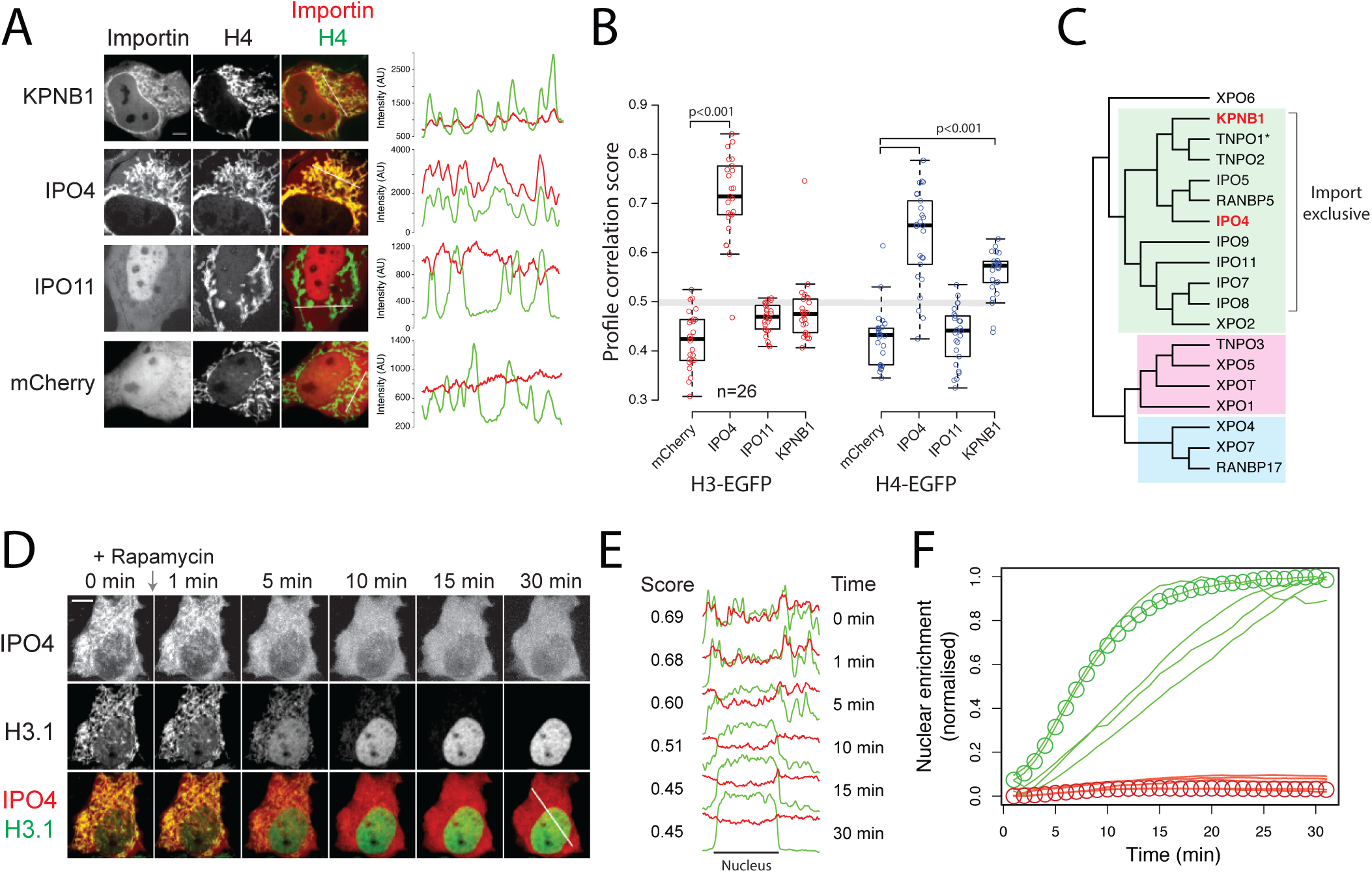
Importing family as cytosolic histone chaperones for monomeric H3.1 and H4. (A) Colocalization between Importin-β proteins and tethered histone H4. Representative images for each importin are shown. Scale bar represents 5 μm. Profiles marked by white lines in the merged channel are shown to the right. (B) Profile correlation analysis between tethered H3.1/H4-EGFP and Importin-β proteins. P-vales of <0.001 from a Wilcoxon rank sum test versus the mCherry alone control are indicated by brackets. (C) Phylogenetic representation of the Importin-β family from humans adapted from (O'Reilly et al., 2011). For further details see the main text. Members found to bind to either H3.1 or H4 in this study are coloured red. An asterisk indicates structural information regarding interaction with H3 (see Figure S4). (D) Time course of H3.1-EGFP release from its mitochondrial tether after co-transfection with mCherry-IPO4. (E) Time course in D shown as a profile that bisects the nucleus and the mitochondrial network. The location of the profile is shown as a white line in the lower right panel of D. Profile correlation scores for each time point are shown. (F) Quantification of nuclear enrichment over time for H3.1-EGFP and mCherry-IPO4 from 5 individual cells. H3.1-EGFP values are represented as green traces, whereas mCherry-IPO4 traces are represented as red traces. Values for the representative cell shown in D is highlight by circles.

Next we asked what happens when the histones are released from their cytosolic tether. To test this we chose IPO4 and H3.1 as an importin-histone pair as they demonstrated strong co-localisation (Figure 5B). H3.1-EGFP^TVMVx2^-FRB-OMP25, mCherry-IPO4 and TagBFP-FKBP12-TVMV-AI were co-transfected into HeLa cells and imaged every minute after rapamycin induced histone release. As H3.1-EGFP was released from its mitochondrial tether, IPO4 diffused away from the mitochondrial network, as expected, but in contrast to H3.1-EGFP, remained cytosolic in its location (Figure 5D-F, Figure S4A, Movie S4). One may have expected a transient increase in nuclear IPO4 levels after H3.1 release as the IPO4-H3.1 complex translocates to the nucleus. However, this would require a sampling time in excess of nuclear import rate. Limited by the kinetics of histone release, our imaging rate is most likely in dearth of nuclear translocation, and thus only the steady state partitioning between nucleus and cytoplasm is observed. Nonetheless, delocalisation of IPO4 from the mitochondrial network and concomitant nuclear accumulation of H3.1-EGFP suggests a hand-off event between the importin and the nuclear histone chaperoning machinery.

Taken together these findings suggest that a number of Importin-β proteins may interact with cytosolically tethered H3.1 and H4 and, in the absence of the core histone chaperoning machinery in the cytosol, supports the idea that Importin-β family members play both a nuclear receptor and histone chaperoning function (Figure 6) (Jakel et al., 2002).

**Figure 6.**
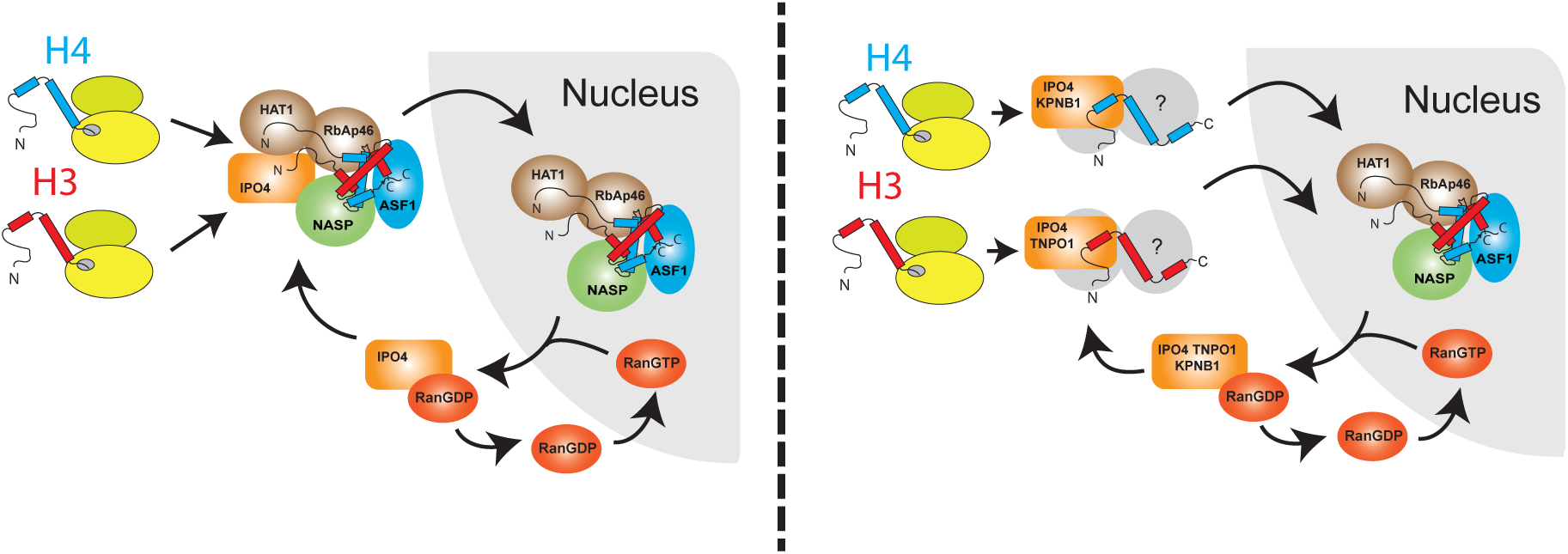
Contrasting models for histone chaperoning during nuclear import of H3.1 and H4. In the model on the left, H3.1 and H4 form a heterodimer soon after synthesis in the cytosol. This heterodimer is bound by the chaperones NASP, ASF1, RbAp46 and HAT1, which further associate with IPO4 before translocation to the nucleus. In the model on the right, H3 and H4 monomers can be bound by a number of redundant Importin-β proteins before translocation to the nucleus, wherein they form a heterodimer and associate with the core histone chaperoning machinery. Additional cytosolic factors may also transiently interact with the monomeric histones.

## Discussion

### Design principles for the RAPID-release approach

To observe dynamics of newly synthesised histones in living cells we needed a system that: (1) enabled labelling of histones in the cytosol, (2) allowed a controlled and expeditious release of the tethered cargo, and (3) was well tolerated by the cell. As histone import is a fast process, we rationalised that sequestration of histones in the cytoplasm soon after synthesis would be necessary to allow time for labelling of the histones (i.e. the folding and maturation of an EGFP tag), and to build up a significant reservoir of labelled protein to observe a pulse-chase. We achieved this by utilising a mitochondrial tail-anchoring peptide (Horie et al., 2002), which had the added benefit of allowing C-terminal tagging, avoiding N-terminal tagging effects (Kimura and Cook, 2001). Although this method worked well, the stability of the tail-anchor insertion necessitated peptide bond cleavage as a means to release the histones from their tether. In addition, we reasoned that gating of the cleavage would be necessary to control the pulse-chase. We considered a number of small-molecule activated protease systems such as the allosteric activation of the cysteine protease domain (CPD) of the *Vibrio cholerae* RTX toxin (Lupardus et al., 2008), and the activation of a split TEV protease through rapamycin-controlled peptide-complementation (Gray et al., 2010;Wehr et al., 2006). However, in the case of the RTX toxin, the small molecule ligand (inositol hexakisphosphate) is present in the cytosol of mammalian cells, and in the case of the split TEV protease, folding after complementation requires tens of minutes to hours for full activation (Gray et al., 2010;Kerppola, 2006;Wehr et al., 2006). Instead we pursued a proximity sensor approach in which an auto-inhibited TVMV protease is both recruited and activated through the addition of rapamycin (Stein and Alexandrov, 2014, 2015). The auto-inhibited TVMV protease was well tolerated by the cell, effectively inactive in its soluble form, but highly active when recruited to its substrate, serving as an excellent switch for the release of the tethered histones.

In comparison to self-labelling tags, such as the SNAP-/HALO-tags, which may influence a target protein's biological function, RAPID-release is compatible with any protein or peptide tag fusion. This may prove beneficial in looking at long term turnover of nuclear proteins as short peptide tags are generally better tolerated than larger globular tags. In addition, the kinetics of RAPID-release are typically an order of magnitude faster that what is achieved with SNAP-tag derived methodologies (Clement et al., 2016;Jansen et al., 2007). However, even with this increase in pulse kinetics, the rate of histone nuclear import was still in excess of the kinetics of histone release. Thus, to effectively model rapid cellular processes, such as the import dynamics of histones, improvements with regards to cleavage kinetics will need to be made. In this regard, the downside to a viral protease's stringent specificity is its lower catalytic turnover, although a reported turnover rate of 11 seconds for TVMV suggests a significant increase in release kinetics is still achievable (Hwang et al., 2000;Nallamsetty et al., 2004;Sun et al., 2010). Despite these drawbacks, the ability to cytosolically tether H3.1 and H4 allowed us to investigate the nucleo-cytoplasmic divide at a level that has previously not been possible, and to observe the incorporation of core histones into chromatin at actively replicating domains in living cells.

### Reassessing the nucleo-cytoplasmic divide within the histone chaperoning pathway

A number of interesting findings came from the ability to tether histones in the cytosol, most notably the stable monomeric nature of the tethered histones and the absence of previously identified cytosolic histone chaperones. The explanation we favour to explain this is that the previously identified histone chaperones are nuclear in living cells, but rapidly leak from the nucleus upon fractionation, giving the impression they are cytosolic when analysed by biochemical means. A previous investigation into the nuclear import of histones put forward the idea that members of the importin-β family may have a dual role in both chaperoning and nuclear localisation of basic proteins as they traverse to the nucleus (Jakel et al., 2002). We probed a number of Importin-β proteins and found IPO4 and KPNB1 associate with cytosolically tethered H3.1 and/or H4, but do not accumulate in the nucleus upon histone release, suggesting a rapid hand-off event to the nuclear chaperoning machinery (Figure 6).

Interestingly, current evidence regarding the binding sites of Importin-β proteins suggest they interact with regions in the histone tails (Baake et al., 2001;Blackwell et al., 2007;Mosammaparast et al., 2002;Soniat et al., 2016;Soniat and Chook, 2016), with a recent crystal structure identifying the H3 epitope that binds to TNPO1 (Soniat and Chook, 2016). A number of soluble nuclear proteins have been shown to bind to histone tails, including HAT1 (H4 residues 5-18) (Wu et al., 2012), Rbap46 (Murzina et al., 2008;Song et al., 2008) & TONSL (H4 residues 12-23) (Saredi et al., 2016), providing a potential hand-off mechanism based on binding site competition between importins and nuclear histone chaperones.

Within the nucleus it is likely that H3.1 and H4 fold rapidly, assuming stoichiometric quantities, however, it also seems plausible that there are provisions for chaperoning monomeric histones. Recently, sNASP has been shown to form a stable complex specifically with monomeric H3 *in vitro,* but not with monomeric H4 (Bowman et al., 2017). In support of this, we found that when forced to be cytosolic, through deletion of its NLS, sNASP interacted with monomeric tethered H3.1, but not to tethered H4. Interestingly, forced cytosolic RbAp46 and HAT1 displayed the opposite binding profile, interacting with monomeric H4, but not H3.1. Further investigation will be required to definitively address whether these chaperones interact with monomeric histones in an endogenous setting. However, such a mechanism would aid the cell in coping with non-stoichiometric increases in soluble histones during periods of replication stress, where DNA synthesis is suddenly halted. Members of canonical protein folding pathways, such as HSP90 and HSC70 may also aid in this process (Campos et al., 2010) or could potentially be involved in excess histone degradation (Cook et al., 2011).

One caveat of our reassessed model is that we cannot explicitly exclude the possibility that H3.1 and H4 rapidly fold with their endogenous histone counterparts in the cytosol after release, but before nuclear import can occur. We think this unlikely, however, as cells outside of S-phase, which undergo limited histone synthesis, followed comparable import kinetics to those inside of S-phase. Furthermore, release of a large quantity of tagged histones is likely to transiently quench the soluble pool of the endogenous counterpart, yet we found nuclear import occurred rapidly, without a soluble accumulation of histones in the cytoplasm.

In summary, our findings suggest that formation of an H3.1-H4 heterodimer is not a prerequisite for nuclear import and that, in contrast to previous models, association with dedicated histone chaperones and folding of the H3.1-H4 heterodimer most likely occurs in the nucleus and not in the cytosol.

## Author Contributions

A. J. B. conceived of the study. A. J. B. and M. J. S jointly carried out the experiments, analysed the data and wrote the manuscript.

## Acknowledgements

We would like to acknowledge Robert Cross, Andrew McAinsh and Anne Straub for access to microscopy resources, Dejana Mokranjac for advice on tail-anchoring proteins, Robert Cross for the monoclonal tubulin antibody and Andreas Ladurner for the polyclonal NASP antibody. This work was supported in full by the Wellcome-Warwick Quantitative Biomedicine Program (Institutional Strategic Support Fund: 105627/Z/14/Z). The authors declare no conflicts of interest.

## Supplementary Material

**>Figure S1.**
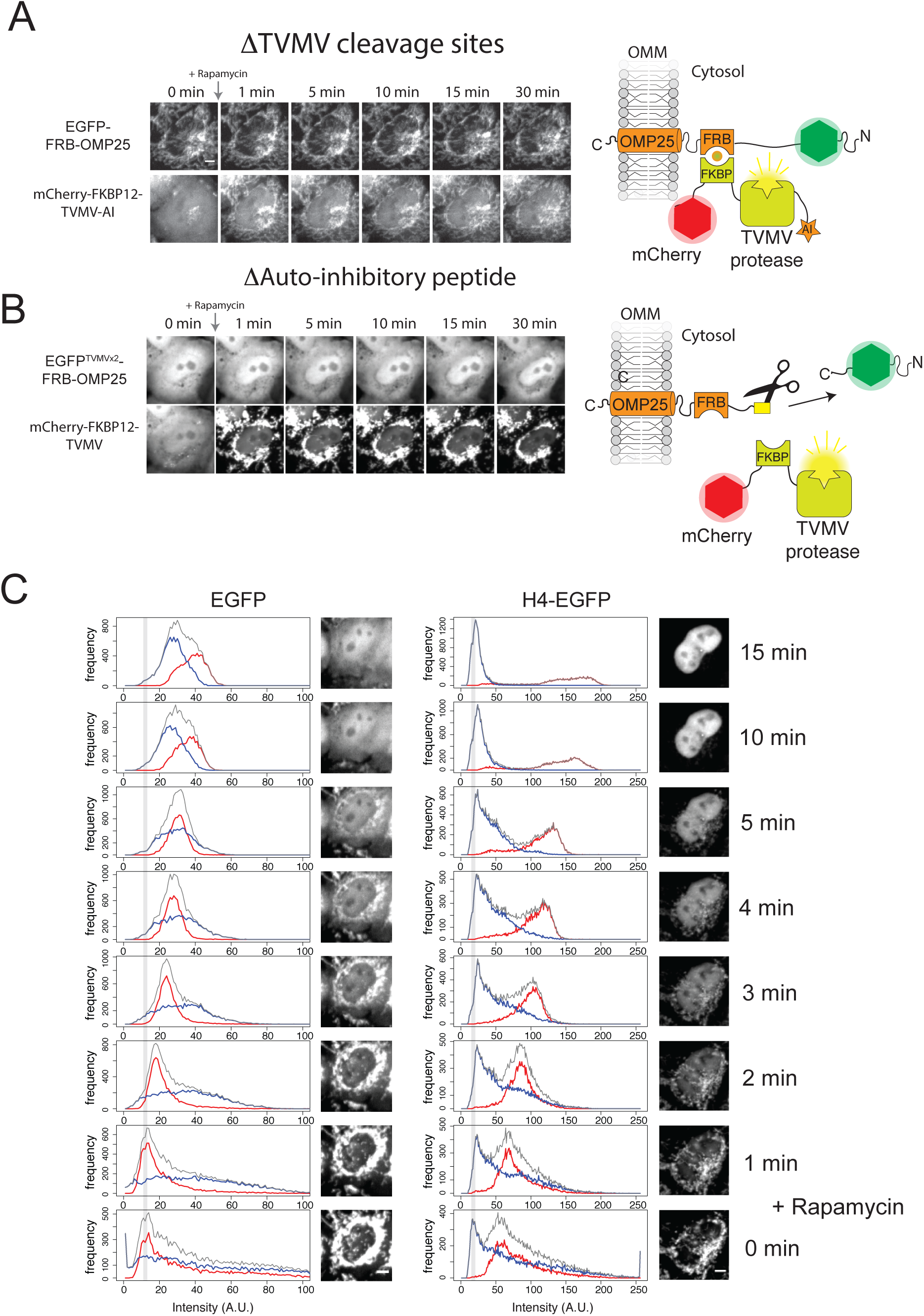
Supplement to Figure 1. (A) Cleavage site deletion mutant. The auto-inhibited TVMV protease is recruited to the tethered EGFP, but is unable to release it due to the absence of a TVMV cleavage site separating EGFP from its tether. Scale bar represents 5 μm. (B) Autoinhibitory (AI) deletion mutant. Deletion of the AI domain from the mCherry-FKBP12-TVMV-AI construct results in constitutively active protease that cleaves the tethered EGFP independent of recruitment. Scale bar represents 5 μm. (C) Histogram of a time course for EGFP or H4-EGFP release. Pixels from the whole cell (grey), the cytosol (blue) and the nucleus (red) plotted for the cells shown in Figure 1E. Whilst the modal value from the whole cell significantly shifts to the right after the release of EGFP, with overlapping cytosol and nuclear signals, the modal value for H4-EGFP remains at a similar level, whilst the nuclear and cytosolic signals form discrete populations. Grey lines represent modal pixel values before rapamycin addition.

**>Figure S2.**
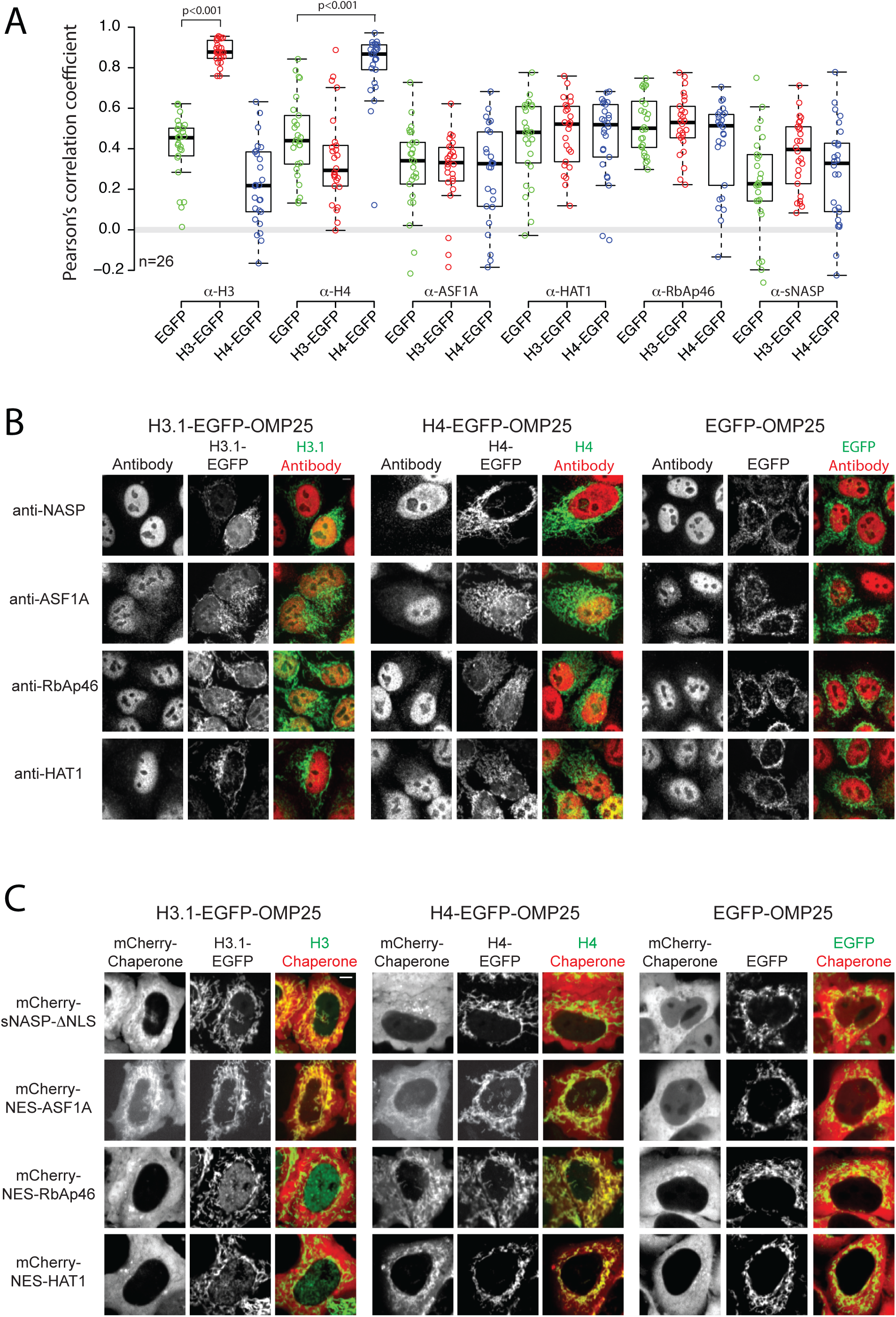
Supplement to Figure 3. (A) Pearson's correlation coefficient for values shown in Figure 3C. P-values of <0.001 from a Wilcoxon rank sum test versus the EGFP alone control are indicated by brackets. (B) Representative images for cells quantified in Figure 3C. (C) Representative images from cells quantified in Figure 3E. Scale bars represent 5 μm.

**>Figure S3.**
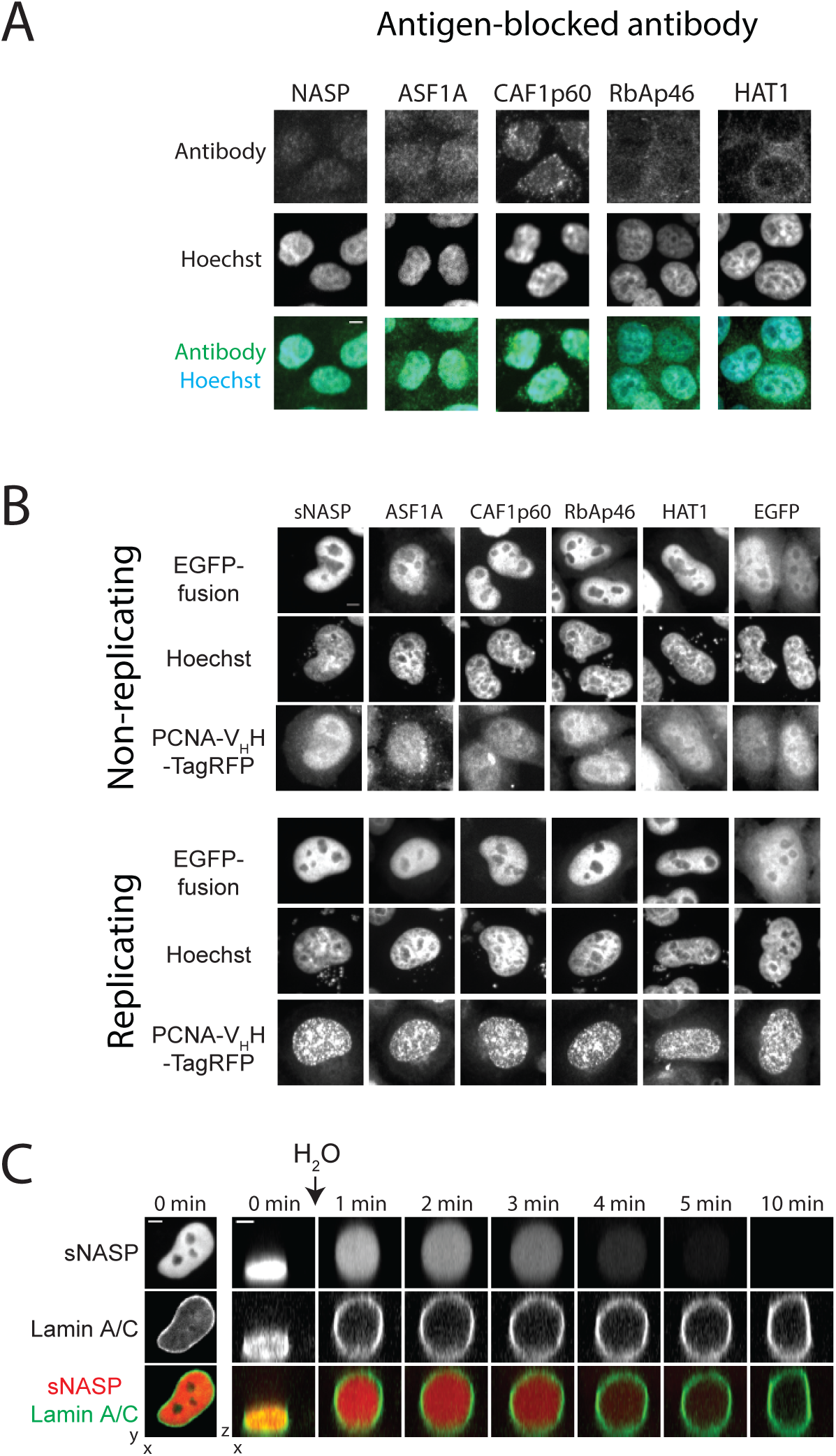
Supplement to Figure 4. (A) Immunofluorescence using antibodies pre-blocked with their immunogen. In each case the specific nuclear fluorescence was decreased to background levels. (B) Cellular location of transiently expressed EGFP-histone chaperone fusions in replicating and non-replicating cells. Scale bars represent 5 μm. (C) Time series of a cell expressing mCherry-sNASP & EGFP-LaminA/C undergoing hypotonic lysis. Scale bar represents 5 μm.

**>Figure S4.**
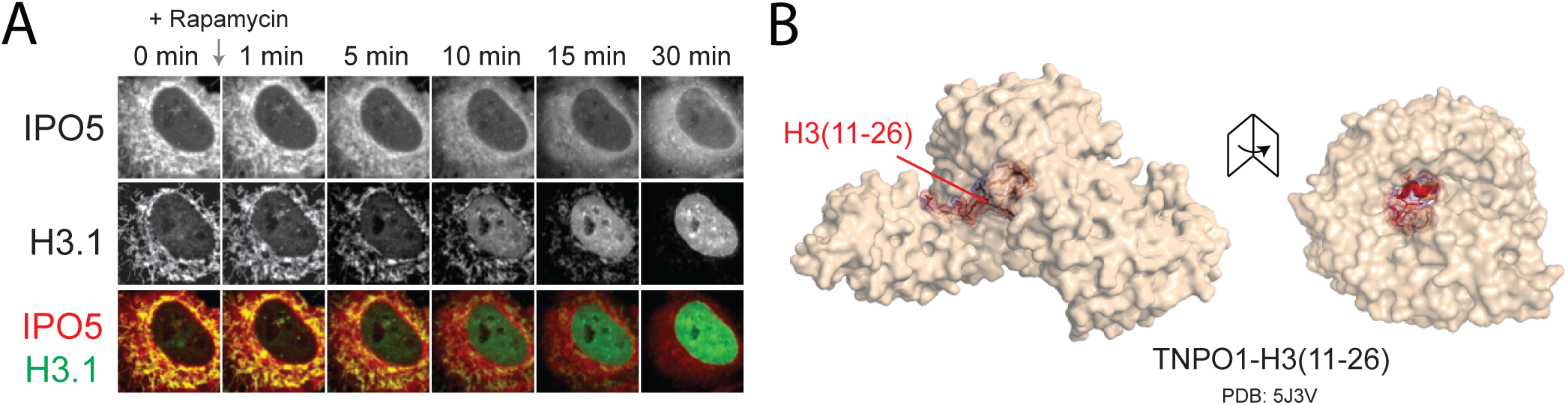
Supplement to Figure 5. (A) A second representative cell showing a time course of H3.1-EGFP release from its mitochondrial tether after co-transfection with mCherry-IPO4. (B) Crystal structure of the closely related TPNO1 bound to a histone tail peptide of H3.

**Movie S1. Related to Figure 1E.** RAPID-release of H3.1-EGFP. Time point ‘0 min' is pre-rapamycin addition.

**Movie S2. Related to Figure 1E.** RAPID-release of H4-EGFP. Time point ‘0 min' is pre-rapamycin addition.

**Movie S3. Related to Figure 4D.** Leakage of mCherry-sNASP from the nucleus upon cell lysis. Time point ‘0 min' is pre-lysis.

**Movie S4.** Related to Figure 5D. Dissociation of IPO4 and H3.1-EGFP during nuclear translocation. Time point ‘0 min’ is pre-rapamycin addition.

## Extended Methods

### Antibodies, plasmids, reagents

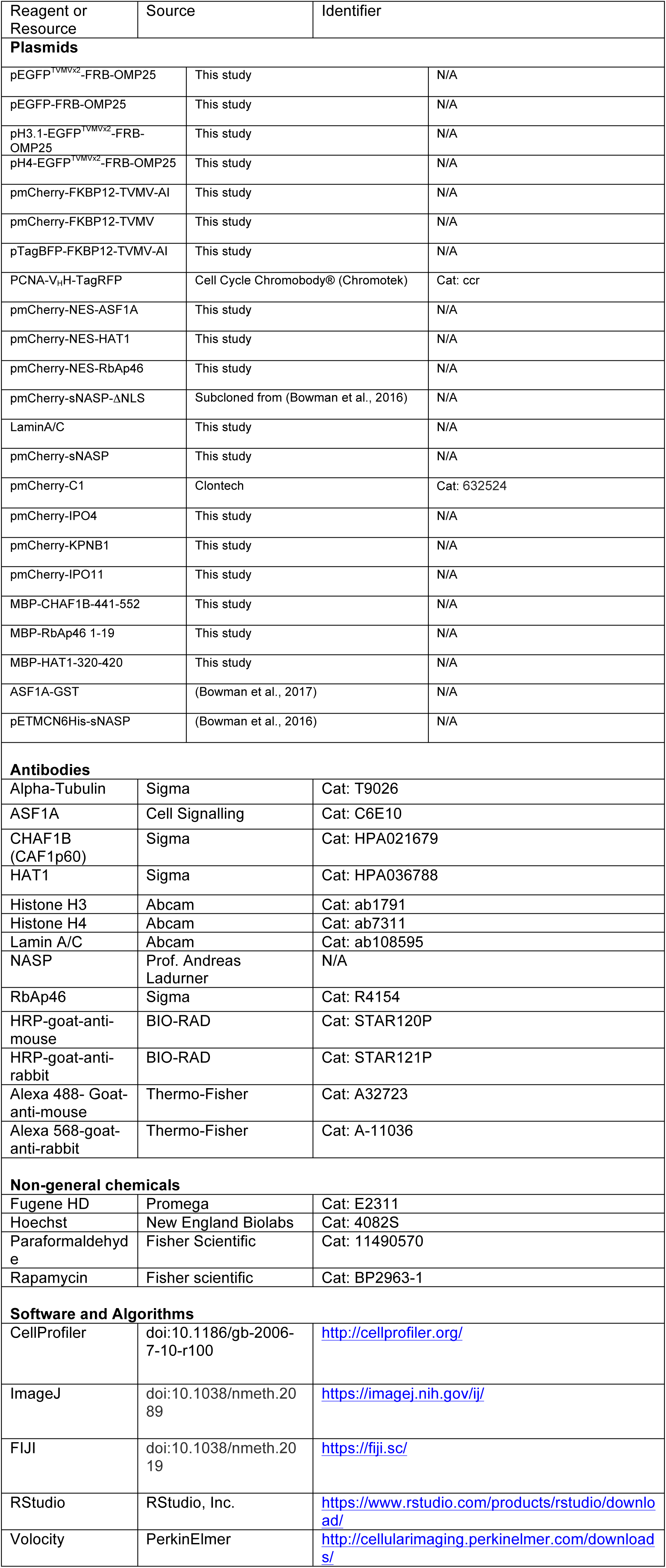

### Cloning and vector assembly

EGFP^TVMVx2^-FRB-OMP25, H4-EGFP^TVMVx2^-FRB-OMP25, H3.1-EGFP^TVMVx2^-FRB-OMP25, EGFP-FRB-OMP25, mCherry-FKBP12-TVMV-AI, mCherry-FKBP12-TVMV and TagBFP-FKBP12-TVMV-AI were constructed through a mixture of PCR cloning, annealed oligo ligation, gBLOCK synthesis (IDT) and Gibson assembly (Gibson et al., 2009)using the mammalian expression vectors pEGFP-C1/C3, pmCherry-C1 and pTagBFP-C1. FKPB12-TVMV-AI and EGFP^TVMVx2^-FRB cassettes were based on published sequences (Stein and Alexandrov, 2014). Open reading frames from IPO4, KPNB1 and IPO11 were amplified from U2OS cDNA and inserted into the HindIII site of pmCherry-C1 using Gibson assembly. EGFP fusions of RbAp46 and CAF1p60 were created by amplifying open reading frames from U2OS cDNA and ligating them into the HindIII-KpnI sites of pEGFP-C1. HAT1 was first cloned into the protein expression vector pETMCN6His using the NdeI-BamHI sites, and then subcloned into pEGFP-C1 using the restriction sites KpnI and BamHI. This transferred the TEV cleavage site from the pETMCN6His vector, extending the linker between HAT1 and EGFP. EGFP-sNASP and EGFP-ASF1A were cloned previously (Bowman et al., 2017; Bowman et al., 2016). mCherry-sNASP-ΔNLS was created by subcloning from a published EGFP-fusion vector (Bowman et al., 2016) into the pmCherry-C1 vector using KpnI and BamHI sites. pmCherry-NES-RbAp46/-HAT1/-ASF1A were created by first inserting a NES sequence into the multiple cloning site of pmCherry-C1 using annealed oligos, followed by subcloning of the chaperones from pEGFP-C1 vectors as described above. The vector encoding PCNA-V_H_H-TagRFP chromobody was purchased from Chromotek Gmbh, Munich. LaminA/C was amplified from a cloned gene and inserted into the vector pEGFP-C1 using Gibson assembly.

### Sequence information for RAPID-release constructs

EGFP^TVMVx2^-FRB-OMP25 (*EGFP,* **<u>TVMVx2</u>, FRB**, *<u>OMP25</u>*):

*MVSKGEELFTGVVPILVELDGDVNGHKFSVSGEGEGDATYGKLTLKFICTTGKLPVPWPTLVTTLTYGVQCFSRYPDH MKQHDFFKSAMPEGYVQERTIFFKDDGNYKTRAEVKFEGDTLVNRIELKGIDFKEDGNILGHKLEYNYNSHNVYIMAD*

*KQKNGIKVNFKIRHNIEDGSVQLADHYQQNTPIGDGPVLLPDNHYLSTQSALSKDPNEKRDHMVLLEFVTAAGITLGM*

DELYKYSDLEGGSGGSGG**<u>ETVRFQS</u>**GGS**<u>ETVRFQS</u>**GGGGSGGSGGAGGGSGGSGG**VAILWHEMWHEGLEEASR**

### LYFGERNVKGMFEVLEPLHAMMERGPQTLKETSFNQAYGRDLMEAQEWCRKYMKSGNVKDLTQAWDLYYHVFR

***RI****GGTGGSGGTGGSGGSGGSGGTGGS<u>ELKLRGDGEPSGVPVAVVLLPVFALTLVAVWAFVRYRKQL</u>*

H3.1-EGFP^TVMVx2^-FRB-OMP25 <u>(H3.1</u>, *EGFP,* **<u>TVMVx2</u>**, **FRB**, *<u>OMP25</u>*):

<u>MARTKQTARKSTGGKAPRKQLATKAARKSAPATGGVKKPHRYRPGTVALREIRRYQKSTELLIRKLPFQRLVREIAQD</u>

<u>FKTDLRFQSSAVMALQEACEAYLVGLFEDTNLCAIHAKRVTIMPKDIQLARRIRGERA</u>GGGSGGGSGGGSGGGALPV

*ATMVSKGEELFTGVVPILVELDGDVNGHKFSVSGEGEGDATYGKLTLKFICTTGKLPVPWPTLVTTLTYGVQCFSRYP*

*DHMKQHDFFKSAMPEGYVQERTIFFKDDGNYKTRAEVKFEGDTLVNRIELKGIDFKEDGNILGHKLEYNYNSHNVYIM*

*ADKQKNGIKVNFKIRHNIEDGSVQLADHYQQNTPIGDGPVLLPDNHYLSTQSALSKDPNEKRDHMVLLEFVTAAGITL*

GMDELYKYSDLEGGSGGSGG**<u>ETVRFQS</u>**GGS**<u>ETVRFQS</u>**GGGGSGGSGGAGGGSGGSGG**VAILWHEMWHEGLEE**

### ASRLYFGERNVKGMFEVLEPLHAMMERGPQTLKETSFNQAYGRDLMEAQEWCRKYMKSGNVKDLTQAWDLYYH

***VFRRI****GGTGGSGGTGGSGGSGGSGGTGGS<u>ELKLRGDGEPSGVPVAVVLLPVFALTLVAVWAFVRYRKQL</u>*

H4-EGFP^tVMVx2^-FRB-OMP25 (H4, *EGFP,* **<u>TVMVx2</u>**, **FRB**, *<u>OMP25</u>*):

<u>MSGRGKGGKGLGKGGAKRHRKVLRDNIQGITKPAIRRLARRGGVKRISGLIYEETRGVLKVFLENVIRDAVTYTEHAK</u>

<u>RKTVTAMDVVYALKRQGRTLYGFGG</u>GGGSGGGSGGGSGGGVATMVSKGEELFTGVVPILVELDGDVNGHKFSVSG

*EGEGDATYGKLTLKFICTTGKLPVPWPTLVTTLTYGVQCFSRYPDHMKQHDFFKSAMPEGYVQERTIFFKDDGNYKT*

*RAEVKFEGDTLVNRIELKGIDFKEDGNILGHKLEYNYNSHNVYIMADKQKNGIKVNFKIRHNIEDGSVQLADHYQQNTP*

IGDGPVLLPDNHYLSTQSALSKDPNEKRDHMVLLEFVTAAGITLGMDELYKYSDLEGGSGGSGG**ETVRFQS**GGS**ETV**

**RFQS**GGGGSGGSGGAGGGSGGSGG**VAILWHEMWHEGLEEASRLYFGERNVKGMFEVLEPLHAMMERGPQTLK**

**ETSFNQAYGRDLMEAQEWCRKYMKSGNVKDLTQAWDLYYHVFRRI**GGTGGSGGTGGSGGSGGSGGTGGS<u>ELKL</u>

*<u>RGDGEPSGVPVA VVLLPVFAL TL VA VWAFVRYRKQL</u>*

EGFP-FRB-OMP25 *(EGFP,* **FRB**, *<u>OMP25</u>*):

*MVSKGEELFTGVVPILVELDGDVNGHKFSVSGEGEGDATYGKLTLKFICTTGKLPVPWPTLVTTLTYGVQCFSRYPDH*

*MKQHDFFKSAMPEGYVQERTIFFKDDGNYKTRAEVKFEGDTLVNRIELKGIDFKEDGNILGHKLEYNYNSHNVYIMAD*

*KQKNGIKVNFKIRHNIEDGSVQLADHYQQNTPIGDGPVLLPDNHYLSTQSALSKDPNEKRDHMVLLEFVTAAGITLGM*

DELYKYSDLEGGSGGSGG**VAILWHEMWHEGLEEASRLYFGERNVKGMFEVLEPLHAMMERGPQTLKETSFNQAY**

**GRDLMEAQEWCRKYMKSGNVKDLTQAWDLYYHVFRRI**GGTGGSGGTGGSGGSGGSGGTGGSELKLRGDGEPS

<u>GVPVA</u> *<u>VVLLPVFAL TL VA VWAFVRYRKQL</u>*

mCherry-FKBP12-TVMV-AI (*mCherry,* **FKBP12**, *<u>TVMV</u>,* **Af**):

*MVSKGEEDNMAIIKEFMRFKVHMEGSVNGHEFEIEGEGEGRPYEGTQTAKLKVTKGGPLPFAWDILSPQFMYGSKA*

*YVKHPADIPDYLKLSFPEGFKWERVMNFEDGGVVTVTQDSSLQDGEFIYKVKLRGTNFPSDGPVMQKKTMGWEASS*

*ERMYPEDGALKGEIKQRLKLKDGGHYDAEVKTTYKAKKPVQLPGAYNVNIKLDITSHNEDYTIVEQYERAEGRHSTGG*

MDELYKSGLRSRAQGGSGGSGG**VQVETISPGDGRTFPKRGQTCVVHYTGMLEDGKKFDSSRDRNKPFKFMLGKQ**

**EVIRGWEEGVAQMSVGQRAKLTISPDYAYGATGHPGIIPPHATLVFDVELLKLE**GGSGGSGGSGGSGGSGGSKALL

*<u>KGVRDFNPISACVCLLENSSDGHSERLFGIGFGPYIIANQHLFRRNNGELTIKTMHGEFKVKNSTQLQMKPVEGRDIIVI</u>*

*<u>KMAKDFPPFPQKLKFRQPTIKDRVCMVSTNFQQKSVSSLVSESSHIVHKEDTSFWQHWITTKDGQCGSPLVSIIDGNI</u>*

*<u>LGIHSLTHTTNGSNYFVEFPEKFVATYLDAADGWCKNWKFNADKISWGSFILWEDAPEDFM</u>SGLVPRGVGR**<u>EYVRF</u>***

**AP**GS

mCherry-FKBP12-TVMV (*mCherry,* **FKBP12**, TVMV):

*MVSKGEEDNMAIIKEFMRFKVHMEGSVNGHEFEIEGEGEGRPYEGTQTAKLKVTKGGPLPFAWDILSPQFMYGSKA*

*YVKHPADIPDYLKLSFPEGFKWERVMNFEDGGVVTVTQDSSLQDGEFIYKVKLRGTNFPSDGPVMQKKTMGWEASS*

*ERMYPEDGALKGEIKQRLKLKDGGHYDAEVKTTYKAKKPVQLPGAYNVNIKLDITSHNEDYTIVEQYERAEGRHSTGG*

MDELYKSGLRSRAQGGSGGSGG**VQVETISPGDGRTFPKRGQTCVVHYTGMLEDGKKFDSSRDRNKPFKFMLGKQ**

**EVIRGWEEGVAQMSVGQRAKLTISPDYAYGATGHPGIIPPHATLVFDVELLKLE**GGSGGSGGSGGSGGSGG<u>SKALL</u>

*<u>KGVRDFNPISACVCLLENSSDGHSERLFGIGFGPYIIANQHLFRRNNGELTIKTMHGEFKVKNSTQLQMKPVEGRDIIVI</u>*

*<u>KMAKDFPPFPQKLKFRQPTIKDRVCMVSTNFQQKSVSSLVSESSHIVHKEDTSFWQHWITTKDGQCGSPLVSIIDGNI</u>*

*<u>LGIHSLTHTTNGSNYFVEFPEKFVATYLDAADGWCKNWKFNADKISWGSFILWEDAPEDFM</u>SG*

TagBFP-FKBP12-TVMV-AI *(TagBFP,* **FKBP12**, *<u>TVMV</u>,* **Af**):

*MSEUKENMHMKLYMEGTVDNHHFKCTSEGEGKPYEGTQTMRIKVVEGGPLPFAFDILATSFLYGSKTFINHTQGIPD*

*FFKQSFPEGFTWERVTTYEDGGVLTATQDTSLQDGCLIYNVKIRGVNFTSNGPVMQKKTLGWEAFTETLYPADGGLE*

*GRNDMALKLVGGSHLIANIKTTYRSKKPAKNLKMPGVYYVDYRLERIKEANNETYVEQHEVAVARYCDLPSKLGHKLN*

SGLRSRAQGGSGGSGG**VQVETISPGDGRTFPKRGQTCVVHYTGMLEDGKKFDSSRDRNKPFKFMLGKQEVIRGW**

**EEGVAQMSVGQRAKLTISPDYAYGATGHPGIIPPHATLVFDVELLKLE**GGSGGSGGSGGSGGSGG<u>SKALLKGVRD</u>

*<u>FNPISACVCLLENSSDGHSERLFGIGFGPYIIANQHLFRRNNGELTIKTMHGEFKVKNSTQLQMKPVEGRDIIVIKMAKD</u>*

*<u>FPPFPQKLKFRQPTIKDRVCMVSTNFQQKSVSSLVSESSHIVHKEDTSFWQHWITTKDGQCGSPLVSIIDGNILGIHSL</u>*

*<u>THTTNGSNYFVEFPEKFVATYLDAADGWCKNWKFNADKISWGSFILWEDAPEDFM</u>SGLVPRGVGR****<u>EYVRFAP</u>****GS*

### Tissue culture and transfection

Hela cells originally obtained from ATCC (HeLa, ATCC^®^ CCL-2™) were expanded and frozen as aliquots in liquid nitrogen as a source stock. Cells were cultured in DMEM (produced in house) supplemented with 10 % heat-inactivated foetal bovine serum (Sigma), 4 mM glutamine and 50 μg/mL penicillin/streptomycin at 37°C in a humidified incubator with 5% CO_2_. Cells were passaged with 0.25% trypsin-EDTA, centrifuged (1000 rpm for 3min) and then resuspended in complete media. Cell counts were performed using a haemocytometer.

Transfections were performed following the Fugene HD manufacturer's instructions (Promega). The day before transfection Hela cells were passaged, counted, and plated at an appropriate density (2.5x10 or 2.5x10 cells per well for a 6 well plate and an 8 well μ-slides (Ibidi), respectively). Transfection mixtures contained 1 μg plasmid DNA in 50 μl DMEM and then vortexing with 3 pl Fugene HD transfection reagent. The transfection mixture was incubated at room temperature for 10 min prior to addition to cells. Cells were transfected with 15 pl or 50 pl of the transfection mixture per well for an 8 well μ-slide or 6 well plate, respectively. Cells were imaged or fixed 24 h post-transfection.

### Imaging

All images were captured using an UltraVIEW VoX Live Cell imaging System (PerkinElmer). Live cell imaging involved culturing HeLa cells in 8 well μ-slides (ibidi) and replacing medium with 200 μl Leibovitz's L-15 medium (ThermoFisher Scientific) supplemented with 4 mM L-glutamine and 50 μg/mL penicillin/streptomycin. For RAPID-release experiments 50 μl Leibovitz's medium containing 1 μM rapamycin was added directly to the well resulting in a final concentration of 250 nM rapamycin. Alternatively, 50 μl 0.5% NP-40 in sterile PBS was added for cell lysis imaging.

For immunofluorescence, HeLa cells cultured on glass coverslips in 6 well plates were fixed with 4 % paraformaldehyde for 10 min. Unreacted formaldehyde was quenched with 50 mM ammonium chloride for 10 min before the cells were permeabilised with 0.2 % triton-X 100 in PBS for 15 min. Cells were blocked in 3 % BSA in PBS for 1 h and subsequently incubated with the primary antibody diluted in 1% BSA in PBS overnight at 4 °C. Excess antibody was removed with successive washes with 0.1% Tween-20 in PBS and then incubated with the appropriate secondary antibody for 1 h at room temperature. After removal of excess secondary antibody cells were stained with 1 μg/ml Hoechst in PBS for 10 mins and then coverslips were mounted using Prolong Gold antifade reagent (ThermoFisher Scientific). Antibody concentrations used were all 1:100 for primary antibodies except anti-NASP (1:5000), and 1:500 for secondary antibodies. For antibody blocking experiments, recombinant immunogens for each antibody were expressed and purified from bacteria (see below) and added to antibodies at a 100-fold molar excess prior to immunostaining.

### Purification of immunogens

6His-sNASP and GST-ASF1A were expressed and purified according to (Bowman et al., 2017;Bowman et al., 2016). The peptide immunogen against which the RbAp46, HAT1 and CHAF1B antibodies were raised (RbAp46 residues 1-19, and HAT1 residues 320-420, CHAF1B residues 441-552, respectively) were expressed as an MBP fusions and purified over Dextrin Sepharose (GE Healthcare) according to manufacturer's instructions. The MBP fusions were then used directly in the blocking experiments without cleavage of the affinity tag.

### RAPID-release methodology and analysis

An initial image stack was taken prior to rapamycin addition, serving as time 0, after which cells were imaged every 1 minute for up to 30 mintues. Partitioning and quantification of the images was carried out using ImageJ. Exponential, logarithmic and linear models were fitted to the data using the statistical analysis software R.

### Nuclear enrichment analysis

CellProfiler was used to analyse images of HeLa cells immunostained for histone chaperones with and without immunogen blocking. Nuclei were segmented using Hoescht staining as a mask, with the 20 pixels surrounding the demarcated nuclei taken as the cytoplasmic region. The mean intensities of the nuclear and cytoplasmic regions were used to calculate the nuclear cytoplasmic ratio.

| Group A | Group B | P-value |
| --- | --- | --- |
| H3.1, 30 min | H3.1, 1 hour | 0.72414553 |
| H3.1, 30 min | H3.1, 2.5.hours | $< 1 \times 10^{-07}$ |
| H3.1, 30 min | H3.1, 5.hours | $< 1 \times 10^{-07}$ |
| H4, 30 min | H4, 1.hr | 0.222567544 |
| H4, 30 min | H4, 2.5.hr | $< 1 \times 10^{-07}$ |
| H4, 30 min | H4, 5.hr | $< 1 \times 10^{-07}$ |

### Co-localisation at sites of active replication

Asynchronously growing cells were transiently co-transfected with H3.1/H4-EGFP^TVMVx2^-FRB-OMP25, TagBFP-FKBP12-TVMV-AI and PCNA-VhH-TagRFP using FugeneHD (Promega), as described above. 24 hours after transfection histones were released by the addition of rapamycin (200 nM) to the cell culture medium at 5 hours, 2.5 hours, 1 hour and 30 minutes before washing with PBS and fixing in 4% PFA in PBS. Cells were imaged using confocal microscopy as described above, taking 10 z-sections that spanned the volume of the nucleus. Cells were manually screened for nuclei that displayed a peripheral pattern of PCNA staining, indicating mid to late S-phase. Due to the large amount of background space outside of the PCNA foci, single Z-slices that bisected the centre of the nucleus were processed to extract masks for the PCNA and histone-EGFP channels using the ImageJ plugin ‘FindFoci' (Herbert et al., 2014). The masks were then used in CDA analysis (Ramirez et al., 2010) using the ImageJ CDA plugin to calculate the Pearson's correlation coefficient (R) for each cell (http://www.sussex.ac.uk/gdsc/intranet/microscopy/imagej). Boxplots of the R-values were created using the program R, and the Wilcoxon rank sum test was carried out to determine significance of the difference in colocalisation over time. P values of less than 0.001 are displayed in Figure 2D. The full list of P-values is as follows.

| EGFP intensity | mCherry intensity | Score |
| --- | --- | --- |
| 21 | 80 | - |
| 25 | 85 | 1 |
| 29 | 89 | 1 |
| 23 | 94 | 0 |
| 22 | 87 | 1 |
| 16 | 99 | 0 |
| 22 | 83 | 0 |

### Co-localisation in the cytosol at the mitochondrial network (Mitochondrial 2-hybrid)

Transfected cells were either imaged 24 hours post transfection, or fixed 24 hours post transfection and stained with the corresponding antibody. A single Z-plane through the cytosol was imaged. Initially Pearson's correlation coefficients between the two channels were calculated for each cell. However, background fluorescence surrounding the nucleus, where the cytoplasm is most dense, coupled to the enrichment of mitochondria in this region resulted in a high level of basal co-localisation (Figure S2A). Although H3.1 and H4 could clearly be differentiated against the background signal using H3 and H4 specific antibodies (serving as positive controls), we pursued an alternative approach in which a correlation score is given dependent on the co-fluctuation of a one-dimensional profile drawn over the mitochondrial network. Where i represents each pixel of a profile, co-increases/decreases of i compared to i-1 for each channel are scored 1, whereas alternate increases/decreases are scored 0. The mean of the scored pixels represents the extent of co-fluctuation and by assumption, co-localisation. A value of 1 represents perfect co-localisation, a value of 0.5 represents no co-localisation, and a value of 0 represents a perfect anti-correlation. An example of the scoring strategy is shown below:

Boxplots of the mean profile correlation scores were created using the program R, and the Wilcoxon rank sum test was carried out to determine significance of the difference in colocalisation over time. P values of less than 0.001 are displayed in corresponding figures. The full list of P-values is as follows:

Figure 3A

| Antibody probe | Group A | Group B | P-value |
| --- | --- | --- | --- |
| H3 | EGFP | H3 | $6.49 \times 10^{-10}$ |
|  | EGFP | H4 | 0.415 |
| H4 | EGFP | H3 | 0.630 |
| | EGFP | H4 | $4.45 \times 10^{-08}$ |
| ASF1A | EGFP | H3 | 0.835 |
|  | EGFP | H4 | 0.763 |
| HAT1 | EGFP | H3 | 0.015 |
|  | EGFP | H4 | 0.453 |
| RbAp46 | EGFP | H3 | 0.068 |
|  | EGFP | H4 | 0.913 |
| NASP | EGFP | H3 | 0.410 |
|  | EGFP | H4 | 0.763 |

Figure 3E

| mCherry probe | Group A | Group B | P-value |
| --- | --- | --- | --- |
| ASF1A-NES | EGFP | H3 | $4.03 \times 10^{-15}$ |
| | EGFP | H4 | $4.03 \times 10^{-15}$ |
| HAT1-NES | EGFP | H3 | 0.098 |
| | EGFP | H4 | $4.03 \times 10^{-15}$ |
| RbAp46-NES | EGFP | H3 | 0.077 |
| | EGFP | H4 | $8.07 \times 10^{-15}$ |
| sNASP-DNLS | EGFP | H3 | $1.15 \times 10^{-10}$ |
|  | EGFP | H4 | 0.949 |

Figure S2

| Antibody probe | Group A | Group B | P-value |
| --- | --- | --- | --- |
| H3 | EGFP | H3 | $6.49 \times 10^{-10}$ |
|  | EGFP | H4 | 0.001 |
| H4 | EGFP | H3 | 0.023 |
| | EGFP | H4 | $9.67 \times 10^{-09}$ |
| ASF1A | EGFP | H3 | 0.696 |
|  | EGFP | H4 | 0.670 |
| HAT1 | EGFP | H3 | 0.592 |
|  | EGFP | H4 | 0.630 |
| RbAp46 | EGFP | H3 | 0.863 |
|  | EGFP | H4 | 0.240 |
| NASP | EGFP | H3 | 0.042 |
|  | EGFP | H4 | 0.580 |

Figure 5B

| Histone tether | Group A | Group B | P-value |
| --- | --- | --- | --- |
| H3 | mCherry | IPO4 | $3.01 \times 10^{-13}$ |
|  | mCherry | IPO11 | 0.004 |
|  | mCherry | KPNB1 | 0.001 |
| H4 | mCherry | IPO4 | $2.33 \times 10^{-10}$ |
|  | mCherry | IPO11 | 0.525 |
| | mCherry | KPNB1 | $1.79 \times 10^{-07}$ |

### Biochemical fractionation of HeLa cells

Cell fractionation was performed following the REAP protocol (Suzuki et al., 2010). Concisely, HeLa cells cultured in a 100 mm culture dish were washed twice with PBS before being scraped off in 1 ml ice cold PBS and collected in a 1.5 ml Eppendorf. The cells were pelleted by centrifugation for 10 sec, the supernatant removed and then the cells resuspended in lysis buffer (0.1% NP-40 in PBS containing protease inhibitors). A whole cell lysate sample was removed before the lysate was briefly centrifuged for 10 sec and the supernatant taken as the cytoplasmic fraction. The pellet was resuspended in lysis buffer, centrifuged for 10 sec, and the supernatant removed. The pellet which contained the nuclear fraction was resuspended in lysis buffer. The cell fractions were diluted in Laemmli sample buffer and boiled for 3 min. Samples of the cellular fractions were run on a 15% acrylamide gel. Proteins were transferred onto a nitrocellulose membrane and then incubated at 4°C overnight with primary antibodies diluted in 5% milk or 3% BSA (all 1:1000 except anti-NASP 1:10000). Membranes were washed with TBST then incubated for 1 h at room temperature with the HRP-conjugated secondary antibody.

### Imaging nuclear leakage during cell lysis

Cells were imaged 24 hours post transfection with mCherry-sNASP and EGFP-LaminA/C. Hoechst at a concentration of 500 ng/ml was added to the cells prior to imaging and incubated for 20 minutes. Culture medium was replaced with PBS immediately prior to imaging. An initial capture of 20 z-stacks penetrating 20 ^m into the culture dish was acquired before addition of NP-40 to 0.1% or replacement of PBS with H_2_O. Z-stacks were then acquired every minute for up to 15 minutes. Orthogonal views were created in ImageJ using the Reslice function. For quantification, Z-stacks were flattened using a maximum intensity projection and the fluorescence as a percentage of maximum for the whole time series was plotted as the mean of normalised values from a total of 15 cells (Figure 4E). The error bars represented the standard error of the mean (s.e.m.).

### RAPID-release of H3.1-EGFP and mCherry-IPO4

RAPID release was carried out as described above. Z-stacks spanning the cell were flattened into a maximum pixel intensity image. The cytosol and nucleus were manually partitioned for each cell and the nuclear enrichment over the cytosol was calculated for each time point. Values for individual cells were normalised between 1 and 0 and plotted on the same axes for comparison.

---

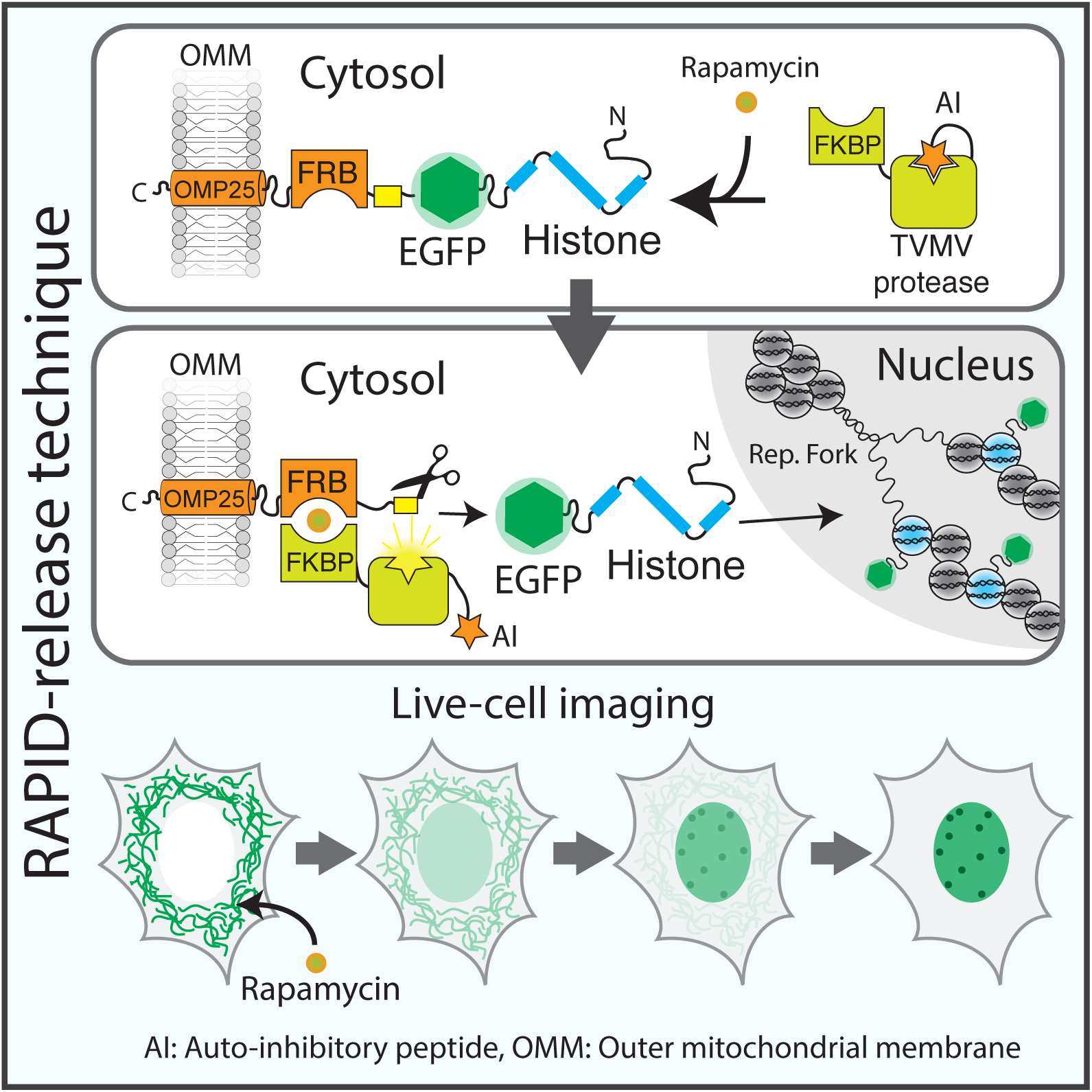

